# Pain as a Trigger for Epigenetic Modifications in Limbic Circuitry

**DOI:** 10.1101/2022.03.29.486268

**Authors:** Svetlana Bryant, Julie-Anne Balouek-Thomert, Luke T. Geiger, David J. Barker, Catherine Jensen Peña

## Abstract

Chronic pain involves both central and peripheral neuronal plasticity that encompasses changes in the brain, spinal cord, and peripheral nociceptors. Within the forebrain, mesocorticolimbic regions associated with emotional regulation have recently been shown to exhibit enduring gene expression changes in models of chronic pain. To better understand how such enduring transcriptional changes might be regulated within brain structures associated with processing of pain or affect, we examined epigenetic modifications associated with active or permissive transcriptional states (histone H3 lysine 4 mono and trimethylation, and histone H3 lysine 27 acetylation) in periaqueductal gray, lateral hypothalamus, nucleus accumbens, and ventral tegmental area five weeks after sciatic nerve injury to model chronic pain. For mice in chronic pain, we observed an overall trend for a reduction of these epigenetic markers in the periaqueductal gray, lateral hypothalamus, and nucleus accumbens, but not the ventral tegmental area. Moreover, we discovered that some epigenetic modifications exhibited changes associated with pain history, while others were associated with individual differences in pain sensitivity. When taken together, these results suggest that chronic pain may lead to a suppression of transcription and gene expression in key limbic brain structures and circuits, which may ultimately result in maladaptive plasticity within these systems.

## Introduction

Chronic pain is a disorder characterized by persistent nociceptive hypersensitivity, often accompanied by allodynia (pain elicited by non-painful stimuli), and hyperalgesia (an amplified response to painful stimuli) (Descalzi et al., 2015). According to the CDC, an estimated 20.4% of the adult population in the United States experienced chronic pain in 2019, and 7.4% of adults were categorized as having high impact chronic pain that limited their lifestyle and work abilities (Zelaya et al., 2020). The limitations of current therapeutics emphasize the need for more effective, safe treatment strategy for chronic pain and a better understanding of its underlying mechanisms.

Pain sensation or perception involve several structures within the nervous system. The dorsal horn of the spinal cord is the initial site for the central pathways of nociceptive information processing. Messages of noxious stimuli are relayed up to the thalamus, parabrachial region, periaqueductal gray (PAG), bulbar reticular formation, and limbic structures (Willis & Westlund, 1997). Beyond traditional pain structures, many other brain structures have been shown to participate in pain processing. For example, manganese-enhanced magnetic resonance imaging in mice with hind paw formalin-induced inflammation revealed changes in the brain activity in the nucleus accumbens (NAc), thalamus, amygdala, and ventral tegmental area (VTA) among other brain regions (Arimura et al., 2019). The ventral tegmental area and nucleus accumbens, structures which play a critical role in reward (Wise & Rompre, 1989) and aversion-associated behaviors (Volman et al., 2013), have also been linked to the affective dimension of pain (Yang & Chang, 2019) and the exacerbation of chronic pain (Gao et al., 2020). In addition, the lateral hypothalamus (LH), which receives primary input from pain-associated structures such as the PAG and habenula, has been demonstrated to be an important part of pain modulation circuitry, with 81% of its neurons responding to noxious stimuli (Dafny et al., 1996).

While acute pain is often treatable, it has become increasingly apparent that chronic pain is a disease of neuroplasticity that requires an understanding of brain structures involved in sensation, emotion, and cognition. Persistent changes in gene expression across brain regions associated with pain (PAG, LH) and with affective and motivational behaviors (LH, NAc, VTA) were recently described (Descalzi et al., 2017; Pryce et al., 2020), implicating enduring epigenetic regulation of gene expression following neve injury. Epigenetic mechanisms may play a role in the maintenance of pain hypersensitivity in rodents undergoing chronic pain (Bai et al., 2010; Cherng et al., 2014; Descalzi et al., 2017; Garriga et al., 2018). However, while previous work has identified a potential role for histone deacetylases in chronic pain at the level of the spinal cord, levels of chromatin modifications associated with active gene expression have not yet been profiled across these supraspinal regions.

Here, we sought to investigate epigenetic modifications associated with chronic pain in C57BL/6J mice subjected to sciatic nerve injury (SNI). We assessed the following epigenetic modifications: tri-methylation on lysine 4 of histone H3 (H3K4me3) and acetylation of lysine 27 of histone H3 (H3K27Ac), both of which are associated with active transcription, and monomethylated H3K4 (H3K4me1), a marker of open chromatin associated with both active transcription or a permissive state that promotes future gene expression. We considered the following key supraspinal structures engaged in pain modulation: periaqueductal gray(Mokhtar & Singh, 2022) and the lateral hypothalamus(Fakhoury et al., 2020). Additionally, we have examined the described epigenetic markers in two structures of the mesolimbic pathway, the VTA and NAc, taking into consideration the growing evidence suggesting an important role of the mesolimbic pathway in chronic pain (Markovic et al., 2021; Watanabe et al., 2018).

## Methods

All Protocols were performed in compliance with the Guide for the Care and Use of Laboratory Animals (NIH, Publication 865–23) and were approved by the Institutional Animal Care and Use Committee, Rutgers University. No animal work was conducted at Princeton University.

### Subjects

Twenty-one C57BL/6J mice (11 males and 10 females, Jackson Labs) were used in the behavioral testing starting at 3-4 months of age. Subjects were group-housed in sets of 3-5 per cage under light/dark 12 h/12 h cycle with access to food and water ad libitum. Mice were randomly assigned to experimental groups SNI (n=11) or sham (n=10).

### SNI Surgery

Mice were anesthetized with isoflurane (1%–4% induction; 1% maintenance) for SNI or sham surgeries. Subsequently, the area from below the knee to the hip on the right leg was shaved and disinfected, an incision along with blunt dissection was made at the mid-thigh level, the muscle layer was separated to expose the sciatic nerve and its three branches. Ligations were then performed on the common peroneal and tibial nerves to segregate a 2-4 mm portion of the nerve that was subsequently removed to generate the spared nerve injury. The sural nerve was left intact. For control mice, a sham procedure was conducted by making an incision at the mid-thigh level and blunt dissecting through the biceps femoris muscle to expose the sciatic nerve. No further perturbations of the sciatic nerve were conducted. Mice were given peri-operative doses of carprofen (5 mg/kg, s.c.) and enrofloxacin (5 mg/kg) and additional dosing for three days following surgery. Subjects were then returned to the colony room for a minimum of two weeks before pain responses were tested.

### Overview of Behavioral testing

Behavioral testing was conducted starting two weeks post-surgery. Different pain tests were executed on separate days. Animals were habituated in the procedure room for 1.5 hours before the beginning of experiments. Female and male subjects were tested separately, animal enclosures and the stand were cleaned thoroughly between experiments.

### von Frey Test of Mechanical hypersensitivity

To evaluate mechanical allodynia, we conducted a manual “ascending stimulus” von Frey test on the subject’s right hind paw. The tested animals were placed in clear animal enclosures with an open floor that was located on top of a grid floor stand. Each von Frey filament of different stiffness (0.08, 0.02, 0.04, 0.07, 0.16, 0.4, 0.6, and 1 gram; Ugo Basile) was applied to the lateral side of the subjects’ hind paws until it bent. When a positive response was recorded, application of the same filament was repeated. Three out of five positive responses indicated the withdrawal threshold for that paw. Five-minute intervals were allowed between the trials.

### Heat hypersensitivity

Heat sensitivity was evaluated using a Hargreaves apparatus (Ugo Basile). Subjects were placed in clear animal enclosures with an open floor on top of a glass stand (Ugo Basile). The infrared source was focused at the center of the paw surface being tested. The infrared source was initially calibrated to elicit a withdrawal time of 10 s from a group of pilot mice that had not undergone any surgical procedures. Animal elicited a withdrawal response, accompanied by flicking or licking of the tested paw. Paw withdrawal time was detected automatically. Five-minute intervals were allowed between each of the three trials.

### Cold allodynia

Cold sensitivity was assessed via the cold plantar assay. Subjects were placed in clear animal enclosures with an open floor on top of a glass stand (Ugo Basile). A cut-off one-milliliter syringe filled with compacted, powdered dry ice was applied to the glass underneath the paw, specifically targeting the edge of the lateral side of the paw surface. The time to paw withdrawal accompanied by flicking or licking of the tested paw was recorded with a stopwatch and documented. Fifteen-minute intervals were allowed between three trials for the same paw.

### Tissue collection assessment

Tissue collection was conducted at 5 weeks post-surgery at least 3 days after the last exposure to pain sensitivity testing, thus reflecting a “baseline” state. Mice were rapidly cervically dislocated, brains were removed rapidly, placed into ice-cold PBS, and sliced into 1 mm-thick coronal sections in a brain matrix. Bilateral punches were made from VTA (16 gauge), lateral hypothalamus (14 gauge), periaqueductal gray (14 gauge), and nucleus accumbens (14G) and flash-frozen in tubes on dry ice.

### Western Blot

Tissue punches were homogenized on ice, in hypotonic lysis buffer (10 mM Tris, 1 mM KCl, 1.5 mM MgCl2, pH 8.0) supplemented with 1 mM Dithiothreitol (DTT), protease inhibitor cocktail (cOmplete™, EDTA-free, Millipore-Sigma) and histone deacetylase (HDAC) inhibitor (10 mM sodium butyrate). Samples were centrifuged for 10 minutes at 10,000g at 4°C to extract nuclei. Supernatants were discarded and pellets were resuspended in RIPA buffer (50 mM Tris, 150 mM sodium chloride, Triton X-100, 0,1% SDS, pH 8.0) supplemented with 10 mM DTT, protease inhibitor cocktail (cOmplete™, EDTA-free, Millipore-Sigma) and HDAC inhibitor (10 mM sodium butyrate). Protein concentration was evaluated with a 660nm Protein Assay (Pierce). Proteins were separated by SDS-PAGE gels (4-20%, BioRad) and transferred to 0.2 μm low-fluorescence PVDF membranes (Amersham Hybond). Proteins were detected with the following antibodies: H3 (1:5,000, ab24834, Abcam), H3K4me1 (1:2,000, ab8895, Abcam), H3K4me3 (1:1,000, ab8580, Abcam), H3K27ac (1:1,000, 07-360, Millipore-Sigma). Secondary antibodies were Alexa Fluor 647 anti-rabbit (1:1,000, 711-605-152, Jackson ImmunoResearch) and AffinipureCy3 anti-mouse (1:1000, 715-165-150, Jackson ImmunoResearch). Images were acquired with Azure Imaging System (Azure Biosystems).

### Statistics

All data are plotted at Mean ± the standard error of the mean (SEM). Western blot results are presented as fold change compared to the sham group and represented as Mean ± SEM. For categorical data, unpaired sample t-tests, two-way mixed ANOVAs, or two-way between-subjects ANOVAs were used, when appropriate. Continuous data were analyzed using multiple regression. Statistics were run using a combination of GraphPad Prism and R. Alpha was always set to 0.05. Post-hoc corrections for the familywise error rate were accomplished using either Sidak or Sidak-Holm corrections.

## Results

### Chronic nature of SNI model

Spared nerve injury is a well-established model of neuropathic pain that induces long-lasting (over 6 months) behavioral modifications (Decosterd & Woolf, 2000). To confirm the chronic nature of pain evoked by the SNI model we conducted weekly behavioral testing on a group of SNI (*n*=5) and sham mice (*n*=5) evaluating their nociceptive sensitivity threshold (**Figure 1A**). We have demonstrated that spared nerve injury produces a robustly lower pain threshold in the SNI mice that persists for at least 5 weeks [**Figure 1B-C**; *von Frey* main effect of Group: *F*(1,16)=14.70, *p*<0.01; *Hargreaves* main effect of group: *F*(1,32)=36.64, *p*<0.001]. To investigate possible sex differences in pain thresholds among male and female mice, we conducted heat and mechanical hypersensitivity tests on male (*n*=32) and female (*n*=29) subjects tested 3-5 weeks following an SNI surgery. Two-way ANOVA revealed a statistically significant difference in average pain threshold when comparing SNI and sham mice in either the von Frey [*F*(1,58)=28.7, *p*<0.001; **Figure 1D**] or Hargreaves [*F*(1,58)= 18.9, *p*<0.001; **Figure 1E**]. However, no differences were observed between sexes in either task.

**Figure 1.**
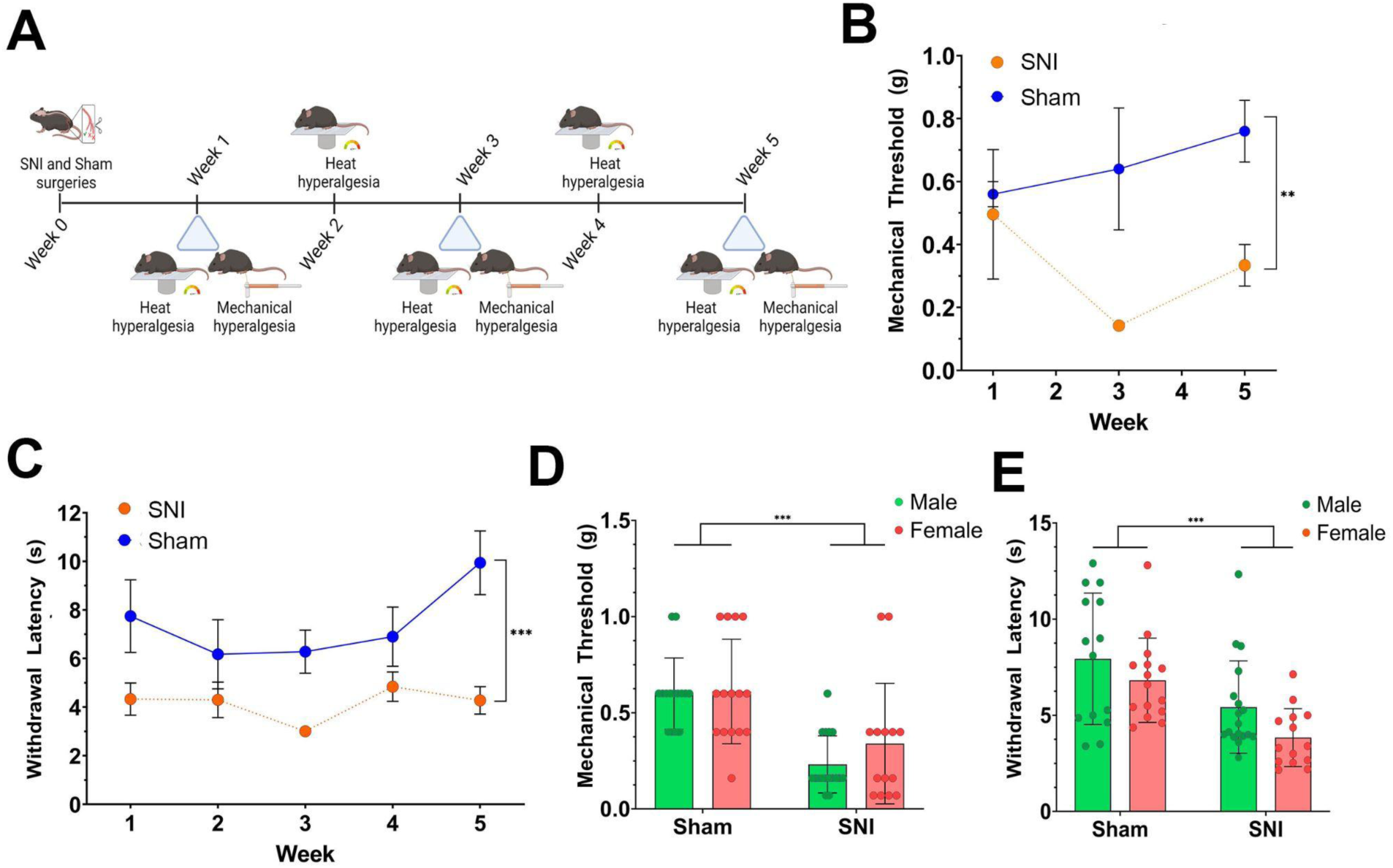
Time course of mechanical and thermal sensitivity for mice with a spared nerve injury. **(A)** Timeline for behavioral testing. Mechanical and thermal withdrawal thresholds were sampled for a period of 5 weeks to determine the optimal point for tissue collection. **(B-C)** Mice with a spared nerve injury exhibited heightened mechanical and thermal sensitivity compared to sham controls (von Frey SNI vs. Sham: F(1,16)=14.70, p < 0.01; Hargreaves SNI vs. Sham: F(1,32)=36.64, p < 0.001). **(D-E)** No sex differences were observed in mechanical or thermal sensitivity following chronic pain (von Frey SNI vs. Sham: F(1,58)=28.7, p < 0.001; Hargreaves SNI vs. Sham: F(1,58)=18.9, p < 0.001)

### Mechanical, Heat, and Cold Pain Sensitivity of C57BL6J Mice Experiencing Chronic Neuropathic Pain

We next examined nociceptive sensitivity thresholds in a group of mice that were used for the investigation of epigenetic markers (**Figure 2A**). One sham control was found to exhibit hypersensitivity to pain and eliminated from further testing. When examining mechanical sensitivity, we found that mice subjected to spared nerve injury (n=11) exhibited significant mechanical hyperalgesia [*n*=10; *t*(19)=6.19, *p*<0.0001; **Figure 2B**], thermal hyperalgesia [*t*(19)=3.39, *p*<0.01; Figure 2C], and cold hyperalgesia [*t*(19)=4.59, *p*<0.001; **Figure 2D**] for their injured paw as compared to sham controls.

**Figure 2.**
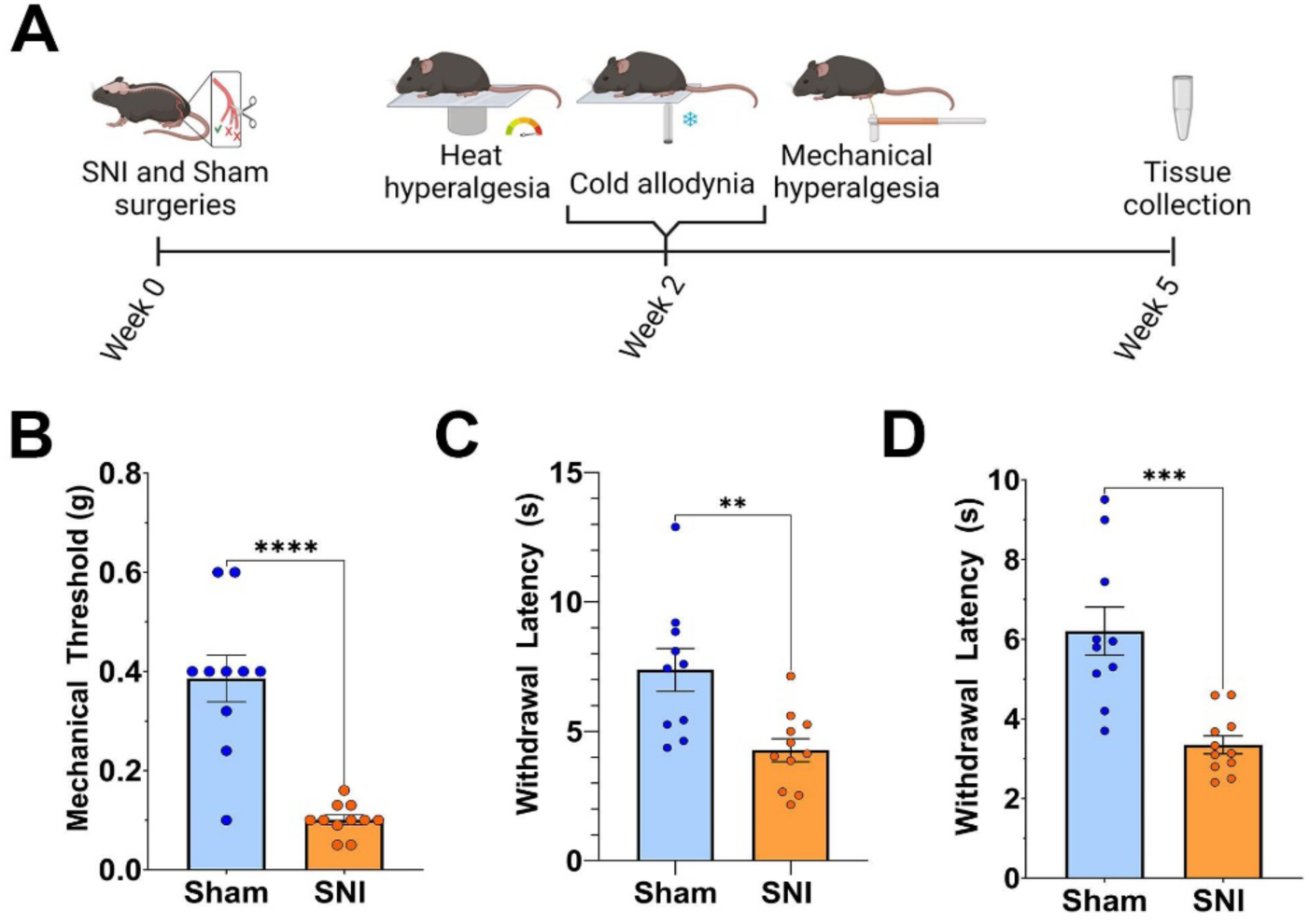
Mechanical and thermal differences in pain sensitivity for mice selected for the evaluation of chromatin modifications. **(A)** Timeline for surgery, testing of pain thresholds, and tissue collection. **(B-D)** SNI mice exhibited hypersensitivity to **(B)** mechanical stimulation (t (19) = 6.19, p < 0.0001), **(C)** irradiant heat (t (19) = 3.39, p < 0.01), and **(D)** noxious cold (t (19)= 4.59, p < 0.001).

### Epigenetic Modifications observed in Mice Experiencing Chronic Neuropathic Pain

We investigated changes in post-translational histone modifications in the PAG, LH, VTA, and NAc by Western blot. In the PAG, we found reduced levels of H3K4me1 [*t*(19)=2.214, *p*<0.05] in mice subjected to spared nerve injury, compared to sham mice (**Figure 3A**). SNI did not trigger any significant change in H3K4me3 nor H3K27Ac levels. Following SNI, H3K27Ac levels were significantly decreased in the LH [*t*(19)=2.731, *p*<0.05] and in the NAc [*t*(18)=3.293, *p*<0.001] compared to sham mice (**Figure 3F,G**). SNI did not affect H3K4me1 nor H3K4me3 levels. In the VTA, we found no effect of SNI on H3K27Ac and H3K4me1 levels. No main effect of sex was found for any of these regions (data not shown).

**Figure 3.**
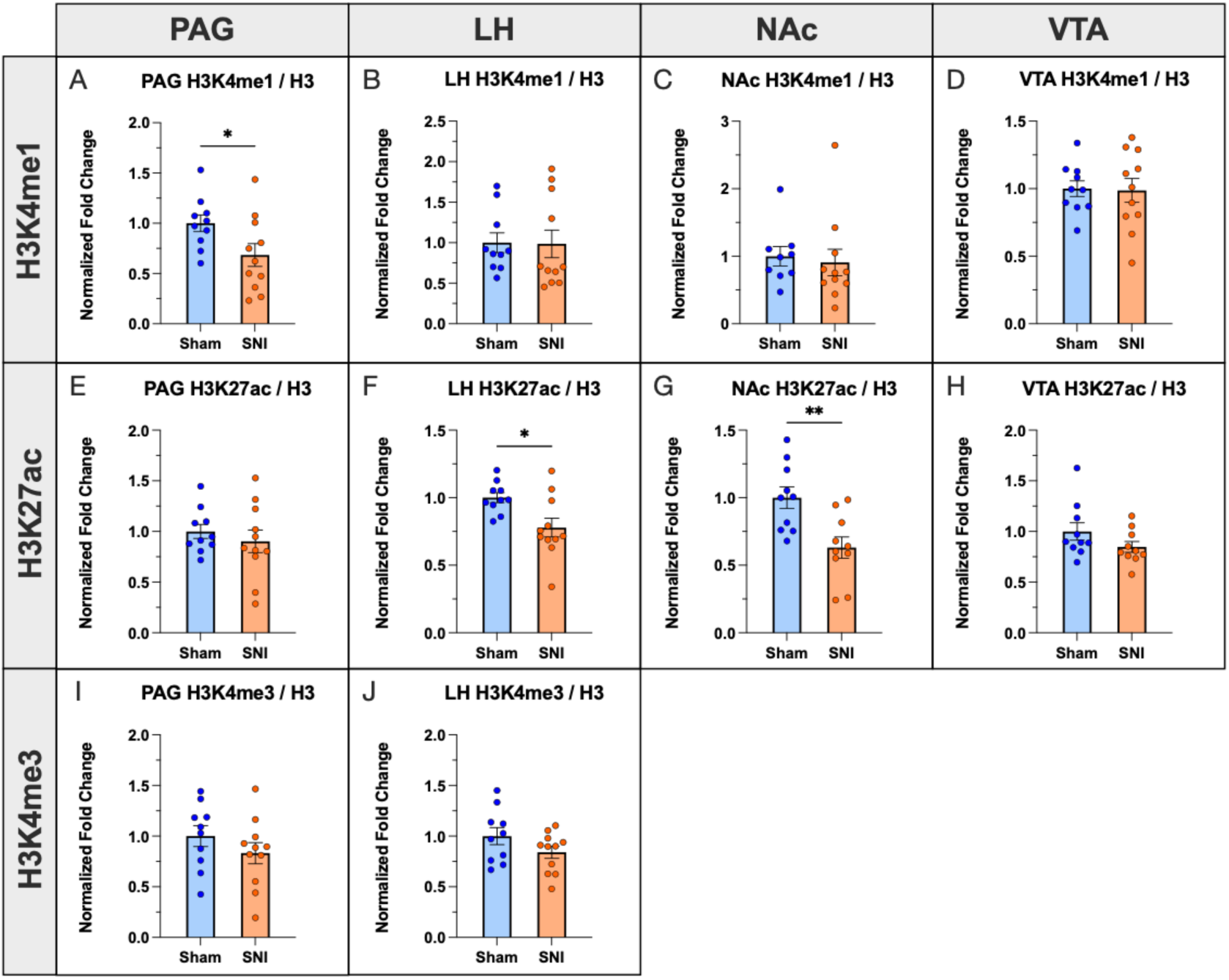
Changes in post-translational histone modifications following spared nerve injury mice. H3K4me1 **(A-D)** was decreased in the PAG in SNI mice compared to sham. SNI reduced H3K27Ac **(E-H)** in the LH and the NAc. No change was found in H3K4me3 levels **(I-J)**.

### Epigenetic changes predict pain severity scores

To determine the relationship between chromatin modifications and pain, we calculated an overall pain *severity* score for all subjects using the summation of their task-specific Z scores (**Figure 4A**). These values were then entered into a multiple-regression analysis to examine the relationship between the pain severity score and the chromatin modifications H3K4me1, H3K4me3, H3K27Ac across the sampled brain regions via multiple regression analysis.

**Figure 4.**
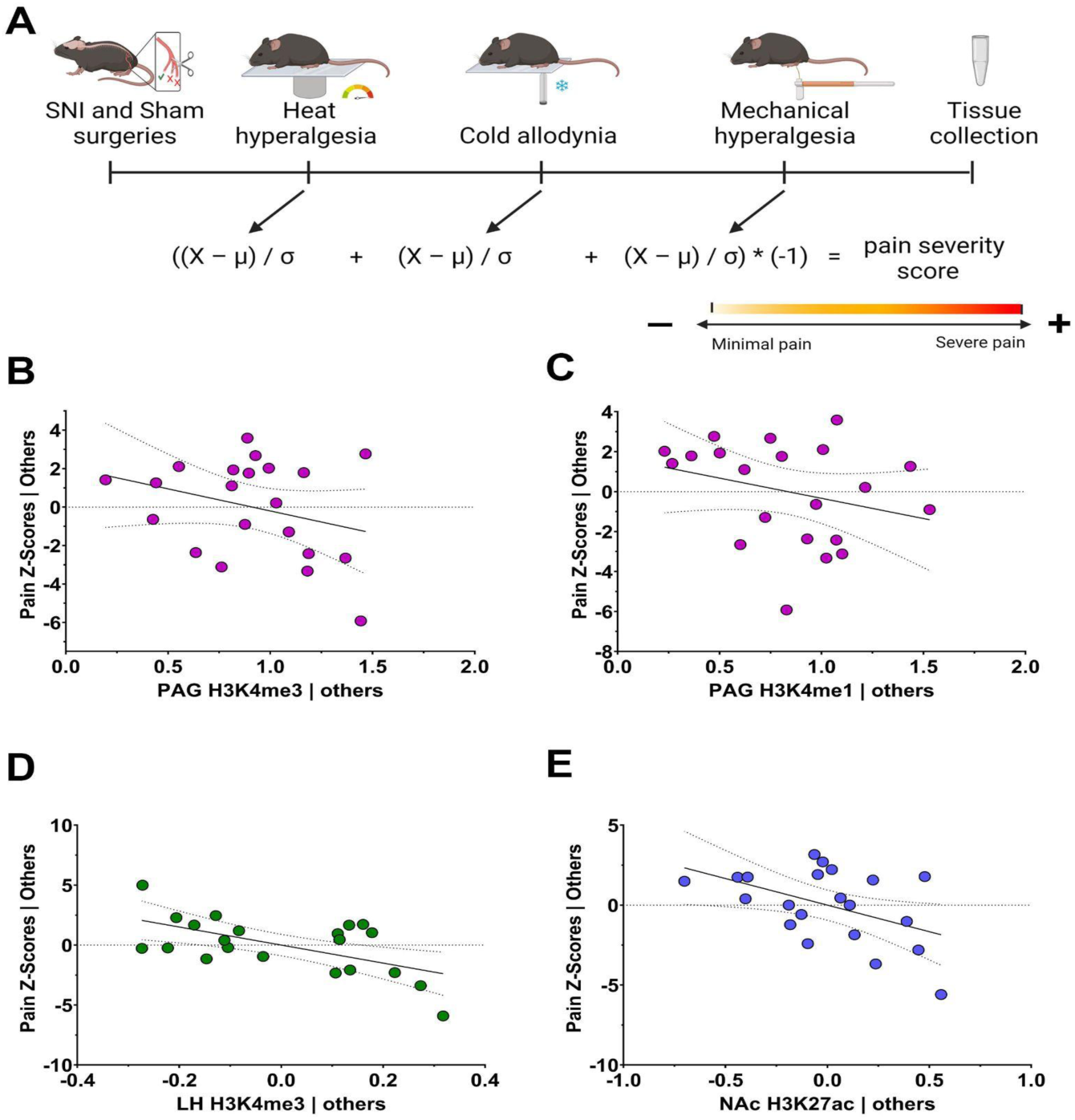
Epigenetic changes predict overall pain severity. (**A**) The z-scores from all pain tasks were taken and combined into an overall pain severity score and correlated with level of each chromatin modification in the PAG (purple), LH (green) or NAc (blue). The data are represented here as partial regression plots, showing the influence of each marker while controlling for the influence of others (**B-E)**. In the PAG, H3K4me3 (**B)** and H3K4me1 (**C**) predicted pain scores. In LH, H3K4me3 (**D)** was associated with pain severity scores. In NAc, only H3K27Ac (**E)** was associated with pain severity score.

Across brain structures, changes in these chromatin modifications in the PAG, LH, and NAc, but not the VTA, were able to predict pain severity scores. In the PAG, levels of both H3K4me1 and H3K4me3 were negatively correlated with pain severity (*R*^2^=0.36, *p*<0.05; **Figure 4 B-C**). Similarly, H3K4me3 in the LH was negatively correlated with pain severity scores (*R*^2^=0.43, *p*<0.05; **Figure 4D**). In the NAc, only H3K27Ac was associated with pain severity scores (*R*^2^=0.36, *p*<0.05; **Figure 4E**)

## Discussion

To better understand how chronic pain drives plasticity in brain structures related to affective processing and pain, we examined epigenetic markers of permissive gene expression state in the PAG, LH, NAc, and VTA. In an SNI model, we found similar behavioral responses for both males and females and observed that hypersensitivity to thermal and mechanical pain was present for at least 5 weeks. At this chronic time point, we also discovered several chromatin changes related to either group differences or individual differences in pain sensitivity, suggesting that chronic pain representations in the brain are encoded by changes in the epigenome.

SNI consistently decreased the chromatin modifications that we examined, with some variability in specific modifications across brain regions. H3K4me1, a marker of open chromatin that is permissive for gene expression — but not necessarily active gene expression — was decreased specifically in the PAG. Because H3K4me1 marks promoters, active enhancers, and enhancers primed to facilitate future gene activity (Calo & Wysocka, 2013; D’Urso & Brickner, 2017), future work should determine whether this decrease in H3K4me1 is specific to enhancers which may indicate limited plasticity in response to future environmental cues rather than reduced active gene expression. In the PAG, the two markers of active gene expression, H3K4me3 and H3K27Ac, were unchanged at the group level, which is consistent with RNA-seq in the PAG after SNI showing equal rates of increased and decreased gene expression (Descalzi et al., 2017). Interestingly however, both PAG H3K4me1 and H3K4me3 predicted individual differences in pain sensitivity, perhaps suggesting that methylation at this residue may serve as a marker of innate differences in pain susceptibility. SNI decreased H3K27Ac in the LH and NAc at the group level, and in the NAc H3K27Ac also predicted individual differences in pain sensitivity. This is consistent with predominantly decreased gene expression found in the NAc 2.5 months after SNI (Descalzi et al., 2017). SNI did not alter these chromatin modifications in VTA, consistent with even levels of increased and decreased gene expression in the VTA after SNI (Pryce et al., 2020). The absence of effects in structures like the VTA with clear roles in pain processing (e.g., Huang et al., 2019) may also suggest that regional heterogeneity or differences in cell-type-specific histone modifications may also be of critical importance. These enduring changes in epigenetic regulation, coupled with the previously identified gene expression changes, may represent either the signature of perceived pain in the supraspinal nervous system or an attempt at adaptive, homeostatic response to pain (i.e., limiting plasticity of gene expression). To distinguish between these possibilities, future work will need to assess epigenetic and gene expression responses after a pain challenge.

Prior investigations into the role of the epigenome in induction and maintenance of pain hypersensitivity have largely focused on histone acetylation in the spinal cord or globally throughout the body. Consistent with our findings of reduced H3K27Ac in the LH and NAc, several studies have found a relationship between reductions in acetylation and pain. For example, spinal nerve ligation (SNL) in rats induced global deacetylation of histone H3 and increased gene expression of the histone deacetylase *Hdac1* in the spinal cord (Cherng et al., 2014), and SNL in mice increased HDAC1 within activated microglia of the dorsal horn of the spinal cord (Kami et al., 2016). Genetic deletion of the histone deacetylase *Hdac5*, resulting in globally increased acetylation, has been shown to increase mechanical pain threshold after SNI (Descalzi et al., 2017). Similarly, intrathecal treatment with HDAC inhibitors has been shown to reduce hindpaw hypersensitivity in rats subjected to various models of neuropathic pain, including antiretroviral drug–induced neuropathy, partial sciatic nerve ligation, L5 spinal nerve transection, or inflammation by complete Freund’s Adjuvant (Bai et al., 2010; Denk et al., 2013). Together, our results and these prior findings are consistent with a general CNS role for histone deacetylation in chronic pain and histone acetylation in pain relief.

Together, our findings and prior work indicate that gene expression may be epigenetically repressed in brain pathways with known roles for pain and reward during chronic pain as a protective mechanism that may also lead to unanticipated, maladaptive rigidity in other contexts. Moreover, we discovered that some markers exhibited notable changes associated with pain history, while others appeared to provide information about individual differences in pain susceptibility.

## Abbreviations

PAG: Periaqueductal gray
NAc: nucleus accumbens
VTA: ventral tegmental area
LH: Lateral hypothalamus
H3K4me1: Histone 3, lysine 4 monomethylation
H3K4me3: Histone 3, lysine 4 trimethylation
H3K27Ac: Histone 3, lysine 27 acetylation
SNI: Spared nerve injury
HDAC: Histone deacetylase

## Acknowledgments

This study was supported by the National Institute on Drug Abuse Grants DA043572 (DJB), the National Institute of Mental Health MH115096 (CJP), and a pilot grant from the New Jersey Alliance for Clinical and Translational Sciences (NJACTs; UL1TR003017 to DJB and CJP). The funders had no role in study design, data collection, data analysis, decision to publish, or manuscript preparation.

## Author contributions

CJP and DJB conceptualized the project and secured funding. SB performed surgeries and analyzed behavioral data. SB and JAB collected tissue for processing and JAB and LTG collected and analyzed western-blot data. All authors contributed to the preparation of figures and writing of the manuscript. All authors have seen the manuscript and approved it for publication.

## Data Availability Statement

All data and analysis code will be made available upon request.

